# When Animals Turn Inside Out: The Eversion of Bloodworms

**DOI:** 10.1101/2025.11.05.686822

**Authors:** Soohwan Kim, Atsingnwi Tuma, David Qin, Young Jae Ryu, Donghui Kim, Aditi Abhilash, Sumedh Chintawar, Caela Thomas-Holness, Arianne Fladger, Essy Behravesh, Ying Zhen, Yanyan Zhou, Joseph T. Thompson, David L. Hu

**Affiliations:** George W. Woodruff School of Mechanical Engineering, Georgia Institute of Technology; School of Biology, Georgia Institute of Technology; Wallace H. Coulter Department of Biomedical Engineering, Georgia Institute of Technology and Emory University School of Medicine; School of Life Sciences, Westlake University; College of Life Sciences, China Jiliang University; Department of Biology, Franklin & Marshall College

**Keywords:** bloodworm, eversion, pressure-driven, hydrostatic skeleton, soft robots

## Abstract

Bloodworms, *Glycera dibranchiata*, possess an eversible proboscis that normally remains concealed within their bodies but explosively everts if the worm attacks or burrows. How does the bloodworm evert quickly and reliably? In a series of experiments, we characterize bloodworm kinematics, pressure, and material properties to estimate the criteria for eversion safely without rupture of the proboscis. We predict the proboscis can withstand pressures 50 times higher and bending strains up to three times higher than the respective values observed. We also present a dimensional analysis of eversion, finding that everting animals, from frogs to snails to sharks, do not satisfy Froude’s law for equivalence of velocities. Our findings may help inspire the development of pressure-driven soft robots with efficient retraction capabilities.

## I INTRODUCTION

To evert is to flip an inside surface outward, as with an eyelid or a sock. For humans, eversion is a medical problem requiring treatment. For example, prolapse occurs when an organ (anus, urethra, or vagina) drops down or bulges out of its normal position due to weakened supporting tissues. However, across the animal kingdom, many animals, including vertebrates and invertebrates, have convergently evolved fast repeatable eversion. Animals evert to burrow, attack, defend, or secrete toxins. In this study, we focus on the bloodworm *Glycera dibranchiata*, one of the many marine worms known as polychaetes that inhabit muddy sediments in such great numbers that they are called “ecosystem engineers” [1, 2]. Many polychaetes evert their pharynx to burrow (Figure 1a), and much attention has been given to the mechanics of the surrounding sediment [3]. The goal of this study is to understand how bloodworms evert by using experiments to relate the bloodworm’s anatomy and material properties to the kinematics and pressure of its proboscis.

**Fig. 1:**
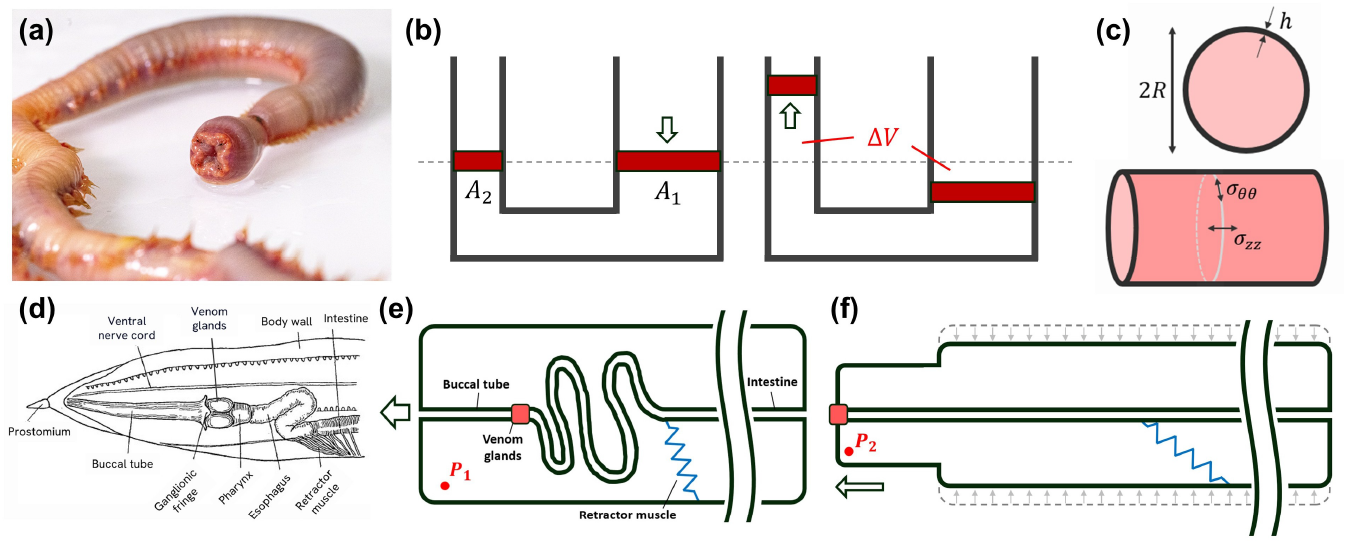
Anatomy and mechanical model of bloodworm eversion. (a) A bloodworm, *Glycera dibranchiata* with an everted proboscis. (b) Piston model for the hydraulic work of the bloodworm, pushing haemolymph of volume Δ*V* into proboscis shown by smaller channel on left. (c) Cylindrical model of the bloodworm’s body with body thickness *h* and radius *R*, where axial (*σ*_*zz*_) and circumferential (*σ*_*θθ*_) stresses are considered. (d) Drawing of bloodworm anatomy, reprinted from Wells [24], highlighting venom glands, buccal tube, and retractor muscles. (e-f) Schematic diagram of the blood-worm’s internal configuration before eversion (e) and after eversion (f), demonstrating extension of the buccal tube.

The biological process of eversion may help provide insights into “vine-inspired” growing robots [4]. Vine robots, like marine worms, extend forward by using pressure to evert internal structures and “grow” new surface area. Thus, there is no sliding or shear force applied to the surrounding environment. One application of vine-inspired robots is in patient transfer, replacing the current method of rolling a patient over to slide a blanket underneath [5, 6]. Challenges in this field include miniaturizing vine robots and ensuring reliable retraction. Various retraction mechanisms have been proposed to address these limitations [7, 8]. Recent advances have enabled the fabrication of vine robots as small as 3 mm in diameter, with potential applications such as medical endoscopy and engine inspection [9]. An understanding of how animals successfully evert and retract may provide inspiration for future designs of vine robots.

We also hope to use this experimental study to improve understanding of hydrostatic organisms, which include blooodworms, snails, caterpillars, and other soft-bodied animals [10]. Even vertebrates can have hydrostatic organs: consider the human tongue, the mammalian penis, or the elephant trunk. Hydrostatic animals (or organs) are often composed of muscles arranged orthogonally to the long axis of the body or organ (e.g., longitudinal, transverse, and circumferential muscle) that enclose an incompressible fluid (e.g., blood, interstitial fluid, cytoplasm). Because the internal fluid is constant in volume, contraction of one group of muscles must cause a change in at least one of the other two dimensions of the animal or organ (e.g., a decrease in body length must be accompanied by an increase in body thickness). Connective tissues are present, often in crossed-fiber helical arrays, that provide structural reinforcement, limit shape changes, and transmit stresses. The relationship between shape and material properties has always been of interest in understanding hydrostatic organisms, and in this study, we shed light on this relationship for bloodworms.

## II Materials & Methods

### A Worm procurement

Since bloodworms are widely used as fishing bait and as a food source for fish and amphibians [11, 12], we obtained bloodworms through bait suppliers (Gulf of Maine Inc., ME, USA). The worms were shipped overnight in insulated styrofoam coolers to preserve freshness. Ice packs were placed at the bottom of the cooler to maintain a low temperature, while wild seaweed was added to provide moisture and cushioning for approximately a dozen bloodworms. The contents were covered with newspaper during transit.

Immediately after receipt, the worms were individually housed in cylindrical plastic pint containers with 200 mL of seawater solution and kept at 10-15^*°*^ C, similar to methods in literature [13]. Artificial seawater with a salinity of 35 ppt (part per thousand) was prepared with Instant Ocean (Spectrum Brands, Blacksburg, VA) and distilled water. The seawater solution was replaced every 2-3 days to reduce the build-up of waste. Worms were housed singly for up to two weeks; during this time, worms were not fed. Total body length in bloodworms was observed to reach up to 35 cm.

### B Worm kinematics

A total of twelve worms were filmed in two media, including air (six worms) and gelatin (six worms). Seawater gelatin simulated the intertidal mud habitat’s mechanical properties [3, 14, 15]. The gel was prepared by mixing 28.35 g gelatin (Sigma-Aldrich, Inc., MO, USA) and 1 L of the same seawater for worm storage. The bloodworms in gelatin were housed in a “sandwich” aquarium constructed of two sheets of acrylic (each 10 mm thick) that were spaced 20 mm apart. Bloodworms were prompted to evert in gelatin by creating a small, shallow hole in the gelatin by pushing a pencil to a depth of a few cm. The anterior end of the worm would then be placed into the hole and the worm would attempt to evert within a minute. Worms in air were placed atop a horizontal plexiglass sheet. The worm’s eversion in air was generally unpredictable, but we elicited eversion by the use of physical contact, salt sprinkles, or patient waiting.

Experiments in air were filmed with iPhone 14 Pro, and in gelatin with the Xcitex-2C camera. The videos were first pre-processed in a video editor (VEED), and the frame rate was downsampled to 10 frames per second. Videos were imported into ImageJ for a frame-by-frame analysis in which automated markers were used to estimate the length and volume of the proboscis. In ImageJ, each frame was converted to 8-bit and then adjusted to apply a threshold to capture only the bloodworm as black. After these settings were applied, the video was cropped to show only the proboscis. The eversion length was measured along the midline of the proboscis at each frame. Then video was analyzed through the built-in function in ImageJ, ‘analyze particles’, which counts the number of black pixels [16] to measure the proboscis area from the top view. Although the proboscis had a baseball bat-like shape, we approximated it as an axisymmetric cylinder with length *L* and radius *R*. We measured the everted proboscis length *L* and the top view area 2*RL*, and estimated the volume as *πR*^2^*L* based on these measurements.

### C Mechanical Characterization

We conducted tensile tests on both worm body wall and proboscis to determine failure strain, failure stress, and Young’s modulus. Four worms were killed by cutting with a scalpel and subsequently bleeding them out. We then cut four samples of body wall and four samples of proboscis. The body wall samples were rectangles with length of 90 mm and width of 25 mm. The proboscis samples were cylinders of 25 mm in length and *D*_*p*_ = 5.5 *±* 0.5 mm in diameter, where the proboscis radius is *R*_*p*_ = *D*_*p*_*/*2. The body wall samples were trimmed into dog-bone shapes, tapering to a width in the middle of *D*_*b*_ = 7 mm. Due to their smaller size, the proboscis samples were left intact without trimming to a dog-bone.

Computing stress from the force requires the cross-sectional area of our samples. The thickness of the sections was estimated from histology: using a body wall thickness of *t*_*b*_ = 770 *±* 190 *µm*, the body wall sample cross-sectional area was *A*_*b*_ = *D*_*b*_*t*_*b*_ = 5.39 *±* 1.33 mm^2^. In the case of the proboscis, modeled as a cylindrical shell, we estimated the cross-sectional area *A*_*p*_ using the measured thickness *t*_*p*_ = 533 *±* 267 *µ*m, resulting in 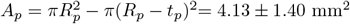.

We performed material testing using an Instron machine (Instron Mark-10) with Bluehill software. We used an elongation speed of 10 mm/s. The instrument was preset to stop once failure was detected, which we defined as a sharp load drop of 400%. To prevent slip of the samples during testing, we blotted dry the clamped portions of the tissue with paper towels. We also affixed sandpaper to the Instron clamps using double-sided tape, which has a higher friction than the smooth clamp surfaces of the Instron machine (Figure 4b).

### D Ultrasound Imaging

We performed ultrasound imaging of the bloodworms immersed in seawater using a Vevo F2 ultrasound system (VisualSonics, Toronto, Canada) and UHF46x transducer (30 MHz center frequency, 46-20 MHz bandwidth), acquiring 2D grayscale ultrasound images at 48 Hz frame rate. Images in the transverse and median sagittal planes were acquired as the transducer was translated using a motion stage (UTS 150 PP, Newport Corp., Irvine, CA) moving down the length of the bloodworm at 0.5 mm/sec. To keep the bloodworm stationary, we designed a 3D-printed open-casket worm “coffin” (Ultimaker 3S) made of a plastic material (PLA), with a width slightly larger than that of the bloodworm. For imaging the everted proboscis, we used ultrasound gel (CanDo, NY, USA, medium viscosity, salt & alcohol free) as the preferred coupling medium. To visualize the non-everted proboscis, the coffin was submerged in artificial seawater, serving as the ultrasound coupling medium. The reason for these two different coupling fluids is that we observed that worms rarely everted underwater, while eversion occurred more reliably in ultrasound gel.

To measure Young’s modulus of the bloodworm tissue, we conducted shear wave elasticity imaging, an ultrasound imaging-based technique for quantification of tissue stiffness [17, 18]. Using the elasticity imaging package on the Vevo F2 and the L38xp transducer (8 MHz center frequency, 10-5 MHz bandwidth; Sonosite, Bothell, WA), we imaged along the transverse and median sagittal planes to acquire the local shear wave group velocity and Young’s modulus.

### E Pressure Measurement

The internal pressure monitoring system was designed to regulate and record intraluminal pressure, where the lumen is the region between the body wall and the intestines. The system, partially adapted from previous work [19], incorporates three pressure transducers (Honeywell SSC series, 1 psi range). One transducer measures the internal pressure approximately 1–3 cm posterior to the fang/venom gland complex, while the other two monitor ambient pressure. These transducers interface with a Texas Instruments (TI) NI Data Acquisition (DAQ) system, which transmits data to a computer via an RS232 serial connection. A custom MATLAB script controls the transducers and logs the pressure data at 0.1-second intervals, allowing real-time monitoring and post hoc analysis of the pressure dynamics.

For pressure measurements, a 25G BD PrecisionGlide needle was mounted onto a probe connected to the pressure transducer. Before each experiment, the pressure transducer was calibrated against the two ambient pressure transducers to ensure consistent baseline measurements. After calibration, the needle was carefully inserted into the lumen of the bloodworm approximately 5 cm posterior to the head. After insertion, pressure was recorded for five minutes for each trial.

We observed clot formation within the needle which impaired pressure transmission. Invertebrates, including bloodworms, possess innate coagulation systems that minimize fluid loss and protect against pathogen entry following injury [20, 21]. Therefore, a new needle was used for each trial.

Simultaneous video and pressure recordings were acquired over five-minute intervals in four trials, each conducted on a different worm. To synchronize kinematic and pressure data, we recorded two simultaneous videos. The first, taken with an iPhone 14 Pro, captured the worm alongside a laptop monitor showing a real-time timestamp (www.timeanddate.com) within the video frame. A second video was filmed of a screen recording of MATLAB’s pressure data acquisition and the same real-time timestamp. The timestamp allowed manual alignment of the onset of eversion and the pressure traces to an accuracy of one second.

## III Mathematical Model

### A Work of eversion

When the bloodworm everts, the work is similar to blowing up a balloon. The total work *W*_*tot*_ may be written as the sum of the work *W*_*p*_ to inflate the balloon, the work *W*_*b*_ to bend the balloon membrane, and *W*_*v*_ the conversion of kinetic energy to heat by viscous action. In total, *W*_*tot*_ = *W*_*p*_ + *W*_*b*_ + *W*_*v*_. We evaluate these terms in turn.

We idealize the worm as a flexible vessel with muscles on the body wall acting as a piston to displace the fluid inside the worm. Then, *W*_*p*_ is the piston work to move hemolymph of volume Δ*V* from the body to its new location within the everted proboscis (Figure 1b). If the fluid pressure during the eversion is *P*, then the work is

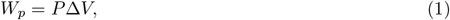

where Δ*V* is the volume of the proboscis, and *P* is the eversion pressure. We write the volume of fluid 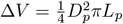 where *D*_*p*_ and *L*_*p*_ are the diameter and length of the proboscis, respectively.

The proboscis must undergo a bending in the process: if we consider a two-dimensional proboscis, by ignoring the larger curvature of the circumference, the work to propagate this fold and evert may be written 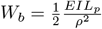 where *E* is the Young’s modulus, *I* is the area moment of inertia, and *ρ* is the radius of curvature. The area moment of inertia 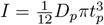 is that of a rectangular beam with thickness *t*_*p*_, and the smallest radius of curvature is *ρ* = *D*_*p*_*/*4 if the beam is folded onto itself as in Figure 5c.

We next consider the viscous dissipation of hemolymph during proboscis displacement. Assuming a parabolic velocity profile across the proboscis radius, the velocity field is given by 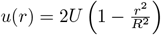 where *r* is the radial distance from the centerline, *R* is the proboscis radius, and *U* is the average flow velocity, taken to be the eversion speed [22]. For unidirectional flow, the local dissipation per unit volume is defined as 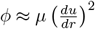. Differentiating the velocity profile gives 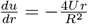 so the squared gradient becomes 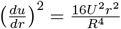. Integrating over infinitesimal annulus of cross-section of 2*πr dr* and length *L*_*p*_, the total dissipation rate becomes 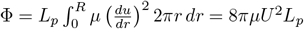. Multiplying by the eversion time 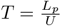, the total work due to viscous dissipation is given by 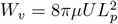.

### B Safety factor

We define two safety factors that describe how closely the worm approaches its material limits. The first safety factor *SF*_*stretch*_ considers unidirectional stretching of the proboscis; the next safety factor *SF*_*bend*_ considers bending as the proboscis is everted from the body.

We define the stretching safety factor *SF*_*stretch*_ as the ratio of the pressure to fracture the proboscis *P*_*frac*_ to the operating pressure required for eversion *P*_*eversion*_:

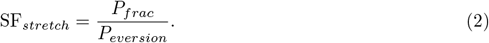

The fracture pressure of the proboscis is calculated using the classical boilermaker equations for a thin-walled cylindrical vessel [23]. One of the boilermaker equations relates the internal pressure *P* to thin-wall longitudinal stress *σ*_*zz*_:

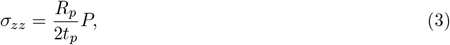

where *R*_*p*_ is the radius of the proboscis (Figure 1c). We neglect consideration of the hoop stress *σ*_*θθ*_ because the material was too small for tensile testing in that direction. Substituting the experimentally measured longitudinal fracture stress *σ*_*zz,f*_ into equation (3) yields the burst pressure 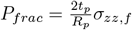 of the proboscis.

The safety factor of bending is written as the ratio of the maximum strain of the proboscis to the strain during 180-degree bending:

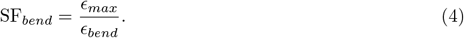

For a bending cantilever, the strain may be written *ϵ*_*bend*_ = *y/ρ* where *y* is the transverse position on the beam and *ρ* is the radius of curvature. At worst case, the radius of curvature is bending upon itself, with *ρ* = *t* and the location of greatest tension is *y* = *t*, yielding *ϵ*_*bend*_ = 1 at the inflection point.

## IV Results

### A Worm Anatomy

We performed histology and ultrasound to determine the bloodworm’s internal structure, which we use to determine the safety factor. Figure 1d shows Wells’ 1937 hand-drawings of a sagittal section of the bloodworm’s head [24, 25]. To identify the same parts in our ultrasonic images, we review the key features drawn by Wells, moving from left to right (anterior to posterior) in the diagram. The proboscis is located posterior to a flap called the prostomium, at the anterior end of the worm. The prostomium plays a role in sensory perception. The buccal tube, resembling a wizard’s hat, is the eversible portion of the proboscis. At the end of the buccal tube is the pharynx. The anterior portion of the pharynx contains the four venom glands and copper-reinforced fangs [26] that secrete neurotoxins [27]. The posterior portion of the pharynx merges with the extensible esophagus. The esophagus connects to the intestine, which then extends posteriorly all the way to the anus. Once the buccal tube is everted from the body, the pharynx and esophagus are enveloped within the buccal tube.

Insets b, c, and d of Figure 2a and Figure 2b-d show closeup views of the venom glands, pharynx, and retractor muscles. Figure 2b shows the cross-section of the worm venom glands and pharynx. The four circular buds at the circumference of the pharynx each contain a fang, a venom gland, and muscles in various orientations. Figure 2c shows a transverse section of the posterior pharynx. Note the relatively thick layer of circumferential and radial muscles that surround the lumen of the pharynx. We presume that this thickened tissue reduces the stress and the chances of fracture during stretch. The outer portions of each circular region contain a layer of longitudinal muscles that run along the entire length of the pharynx (Figure 2c) and continue along the outer portion of the esophagus as shown by the 23 numbered longitudinal muscles in Figure 5d. The histology also revealed that the thickness of the body wall and the buccal tube are *t*_*b*_ = 770 *±* 190*µ*m and *t*_*p*_ = 533 *±* 267*µ*m, respectively (Figure 5d).

**Fig. 2:**
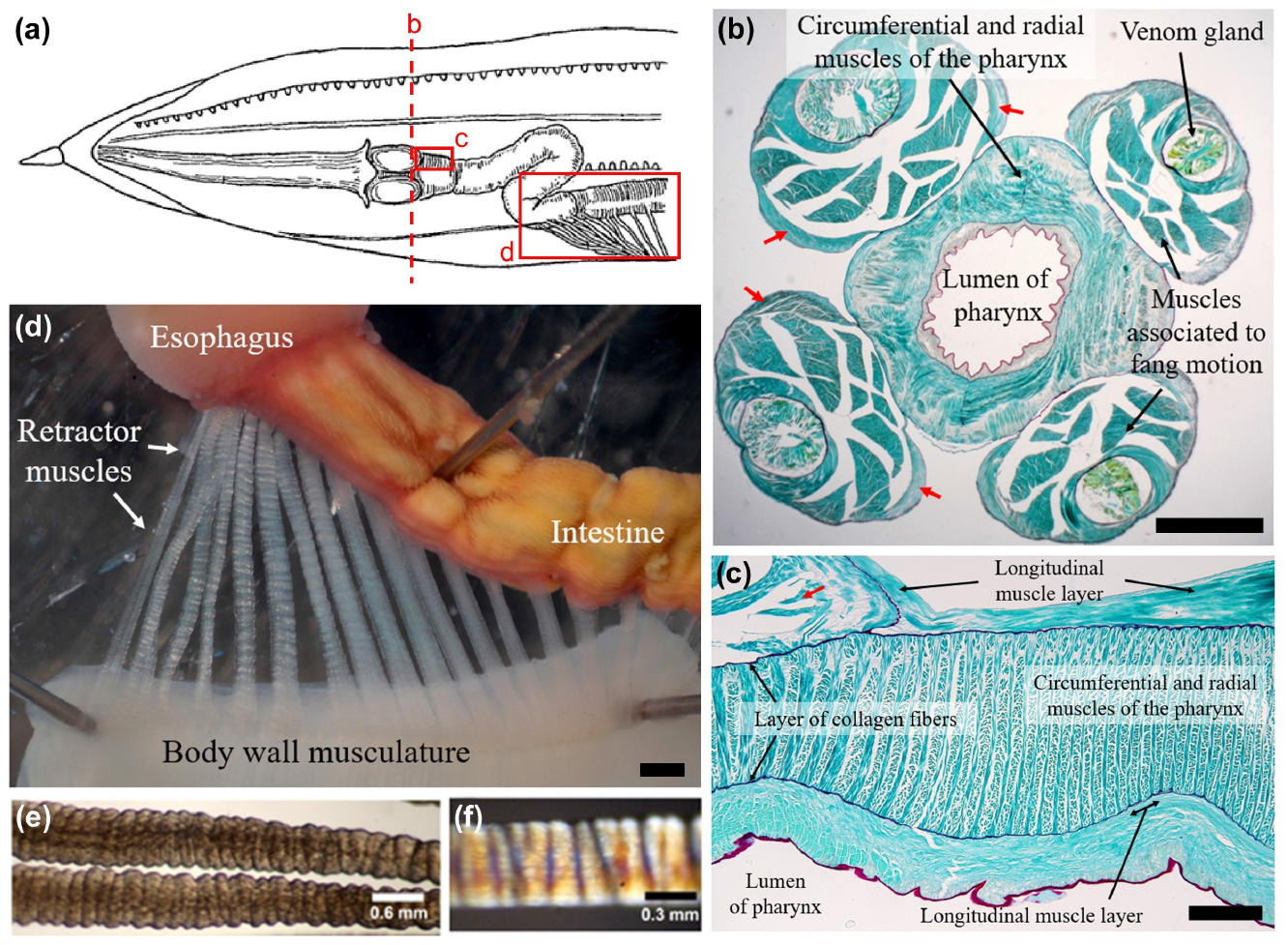
Histological analysis of bloodworm structures. (a) Schematic diagram of the bloodworm showing locations for histological and dissection analyses, reprinted from Wells (1937)[24]. (b) Transverse histological section of the pharynx near the fang/venom gland complex. Red arrows denote the longitudinal muscle layers. (c) Longitudinal histological section of the upper portion of the pharynx. The red arrow indicates the base of the fang/venom gland complex. (d) Dissection of the posterior esophagus, the anterior portion of the intestine, and approximately 17 retractor muscles connecting to the body wall musculature. (e) Transmitted light photomicrograph of moderately contracted retractor muscles. Note the folding. (f) Photomicrograph of a moderately contracted retractor muscle taken under crossed polarizing filters to highlight the banding pattern. Photomicrographs were reprinted from Taylor (2025)[77].

In Figure 2d, the dissection illustrates the posterior end of the esophagus, the anterior portion of the intestine, and approximately 17 mesentery muscles, referred to as “retractors” by Wells. The retractor muscles facilitate the retraction of the proboscis after eversion. They originate on the dorsal midline of the body wall musculature and insert on the intestine and posterior portion of the esophagus. As the buccal tube is everted, the retractor muscles are stretched several times their length. To prevent failure, they are highly wrinkled as shown in the photomicrographic views (Figure 2e,f). This wrinkling allows for a high strain without failure.

Figures 3a,e show ultrasonic imaging of the sagittal view of a worms in its resting and everted state, respectively. This unique view into a living worm revealed several adaptations that accommodate the proboscis inside the worm’s body. In the worm’s resting state, the buccal tube and esophagus are both buckled, showing even more buckling than Wells’ original drawings (Figure 1d), which only includes the esophagus as buckled. When deflated and inside the body, the buccal tube is highly compact, showing a diameter of 2 mm, which is 60 percent smaller than the inflated diameter (5 mm) shown in the everted ultrasonic image in Figure 3e.

**Fig. 3:**
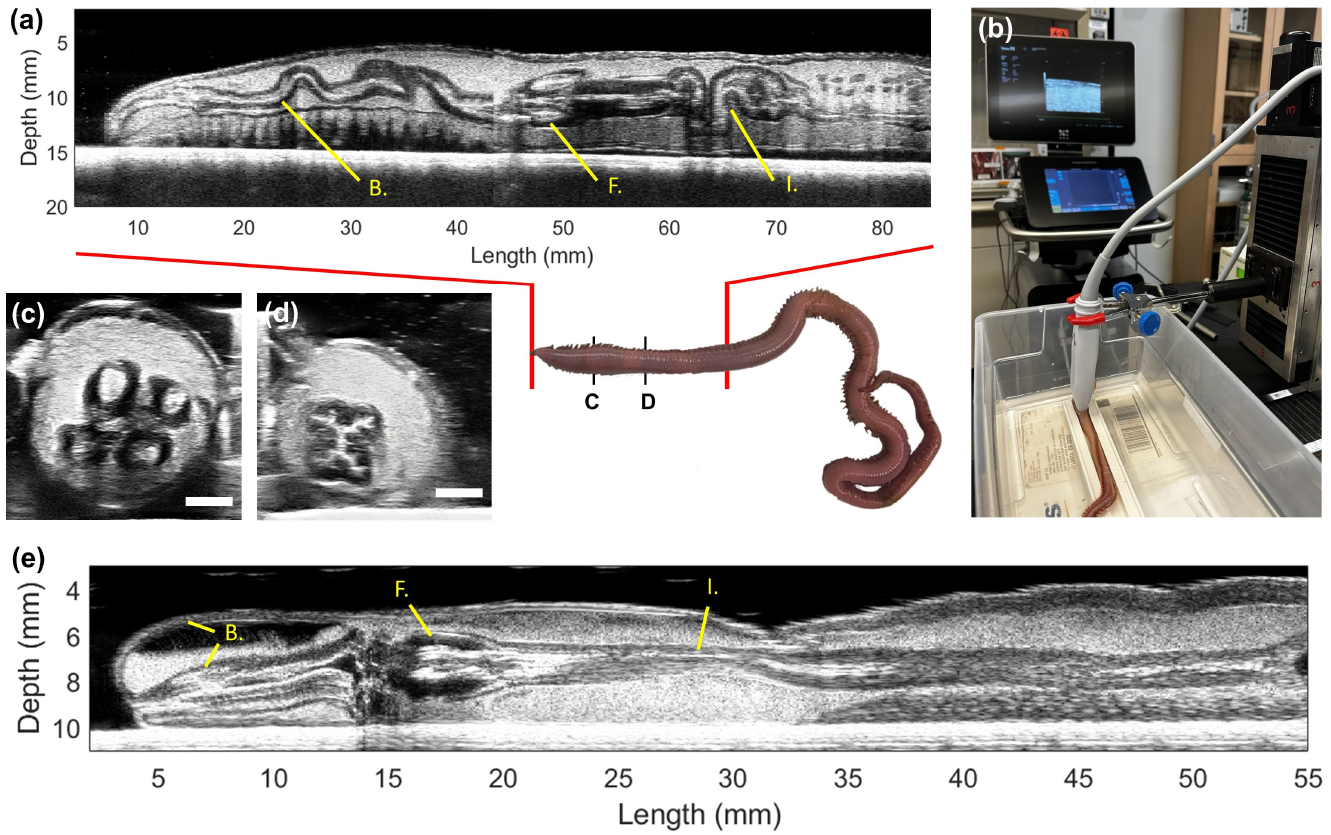
Ultrasound imaging of bloodworm. (a) Sagittal view of worm at rest, with proboscis inside body. This image is composed of two horizontal images stitched together at length of 42 mm. B stands for the buccal tube, F is the fang/venom gland complex, and I represents the intestine. Setup for ultrasound scanning of a bloodworm. (c) Transverse image of the fang/venom gland complex. (d) Transverse image of the buccal tube. Scale bars, 2 mm. (e) Sagittal view of an everted worm.

The everted worm was bathed in ultrasound gel, which we found changed their behavior. Specifically, they no longer fully retracted their proboscis, causing the worm in Figure 3d to be in a half-everted state. After 5 minutes in this state, hemolymph settled to the bottom of the buccal tube, as shown by the white region in Figure 3e. The hemolymph appears to be actively pumping, as shown by the motion of the speckle pattern in Supplementary Videos 1-2. Figures 3c,d show transverse views from the cross sections marked in the adjoining schematic. Figure 3c shows the muscular structure of the venom sacs and Figure 3d shows the wrinkled structure of the buccal tube, which we speculate is an example of self-organized origami [28] to reduce the space taken by the buccal tube when inside the body.

### B Material testing

Having established the anatomical structures inside the worm, we proceed with material testing. We performed tensile testing with four samples each of the worm body wall and the proboscis. Figure 4a shows the stress-strain curves for both materials, the body wall in black and proboscis in red. In the linear regime, the proboscis with a Young’s modulus of 28.6 *±* 14.5 kPa, is half as stiff as the body wall with a Young’s modulus of 73.71 *±* 24.15 kPa. As shown in Figure 4d, the proboscis has stiffness comparable to human muscle [29] while the body wall has stiffness comparable to that of human tendon. A stiff body wall makes sense because it protects the worm from damage on an everyday basis. During ultrasound imaging, we also conducted shear wave elastography [17] to estimate the elastic modulus of the bloodworm’s body. Across four measurements, we obtained an average modulus of 36.2 *±* 11.7 kPa, which is on the same order of magnitude as those obtained from the tensile tests. The elastography measurements were not localized to specific anatomical regions.

**Fig. 4:**
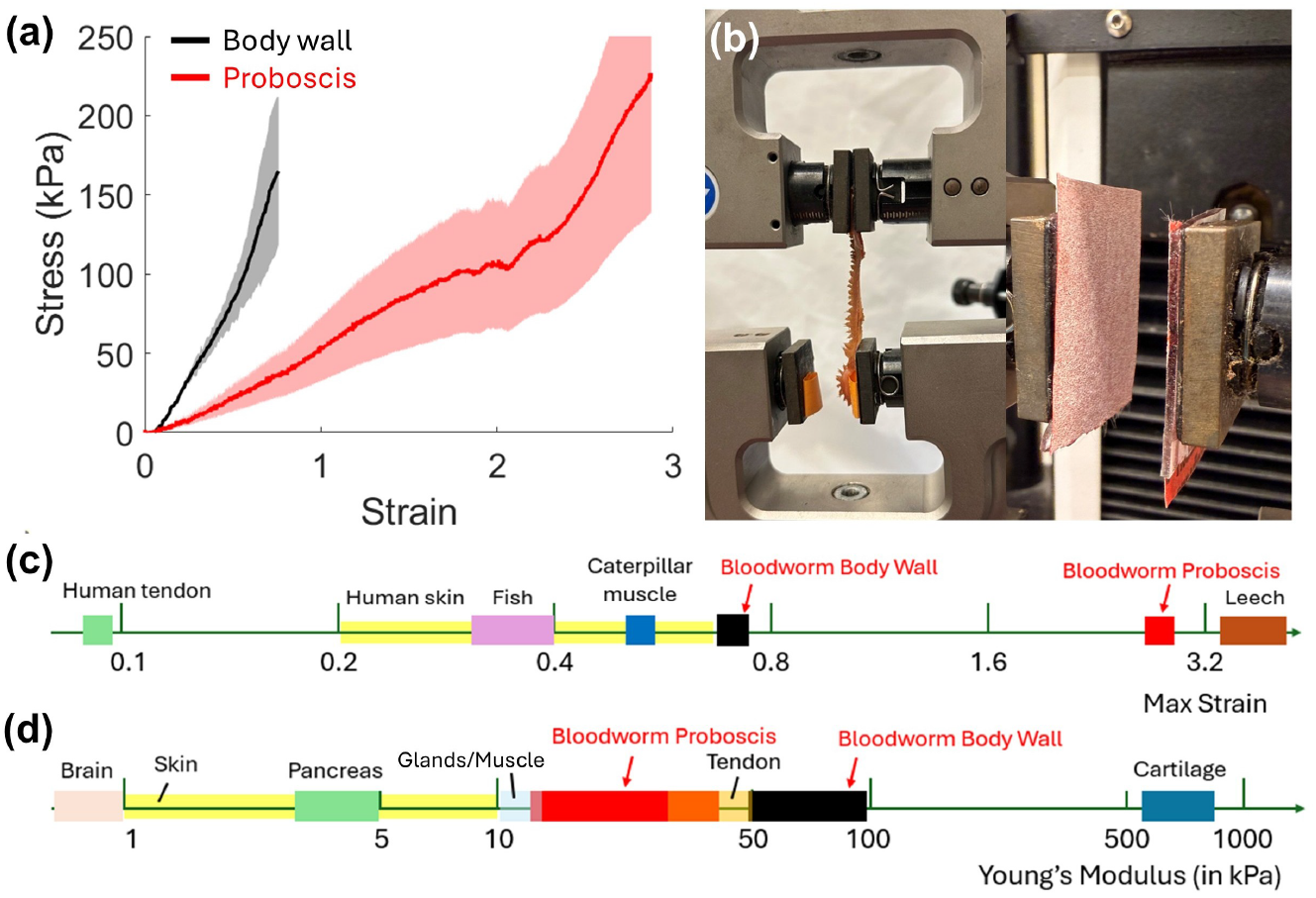
Mechanical characterization of bloodworm tissues using tensile testing (a) Stress–strain curve of the bloodworm body wall (black) and the proboscis tissue (red). (b) Tensile testing setup with a body wall sample in an Instron mechanical tester. Sand paper was used to enhance the grip. Maximum strain of biological tissues with bloodworm proboscis and body wall in red and black respectively. (d) Young’s modulus of biological tissues.

We stretched the materials to failure and found that the proboscis and body wall have a maximum strain of 2.9 and 0.76, respectively. Not only is the proboscis half as stiff as the body wall, but it can also stretch by a factor of three without breaking. Its maximum strain of *ϵ*_*max*_ = 2.9 is comparable to one of the most extensible soft tissues in nature, the body of the leech *Hirudo nipponia* [30] with a max strain of 3.6. In comparison, human tendons fail at a strain of 0.098 [31]. Fish skins are less ductile with a rupture strain of up to 0.3-0.4, but much higher strength than the worms with a rupture strength of 10-15 MPa [32] which is nearly 50 times that of the bloodworm proboscis (*σ*_*zz,f*_ = 230 kPa).

### C Kinematics

We proceed with using videography and pressure probes to measure the dynamics of eversion. Bloodworms are residents of the intertidal mudflats, so they have opportunities to evert in both air and mud. Thus, we performed experiments in both air and gelatin, which mimics mud. Figure 6a shows a time series of bloodworm eversion and retraction in air. Moving left to right in this image sequence: the prostomium flap opens up and the proboscis emerges as a sharp tip. As the buccal tube emerges further, it widens, appearing as a cylinder. The eversion ends with the proboscis as a club shape. The fangs emerge during some eversions, but most often the eversion stops and reverses direction before this point. In eversion, the widest “club” part of the proboscis emerges last. However, the retraction sequence is not simply the time-reversal of eversion. For instance, the base of the proboscis retracts first in retraction so that the fangs stay exposed for as long as possible. This makes sense as the retraction of a prey item requires that fangs bite the prey firmly. We hypothesize that the worm uses both the retractor muscles attached to the dorsal midline of the body wall and the longitudinal muscles in the proboscis for retraction. Both eversion and retraction may require coordination of these muscle groups to retract the base of the proboscis first.

Figure 6b shows the time course of the proboscis length for a representative eversion (Supplementary Video 3). The proboscis reaches a peak length of approximately 50 mm within four seconds (drawn in blue), followed by a brief waiting phase with its fangs out (green) and a slower 8-second retraction (red). Hollow circles correspond to the time points of the images in Figure 6a. The fangs became visible at the onset of the wait phase and disappeared after approximately 80% of the proboscis had retracted. The fast eversion likely evolved to attack prey before the prey can escape, enabling the worm to complete full extension within 1.7 *±* 0.8 seconds. We noticed that worms had two modes of eversion, a fast eversion at 33.8 *±* 4.0 mm/s and a slow eversion at 8.6 *±* 3.2 mm/s. We did not notice any intermediate speeds, suggesting that eversion for worms is like flipping a switch, without much autonomous control.

Figures 6d,e show the proboscis displacement for six trials in air and gel, respectively. Gelatin provides a greater resistance to penetration with eversion in gelatin occurring at speeds of 3.4 *±* 1.5 mm/s, which is ten times slower than the fastest eversions in air (Figure 6c). Eversion through gelatin is associated with a maximum proboscis length of 20 mm, which is less than half value in air of 50 mm (Supplementary Video 4).

### D Pressure testing

Having established repeatable kinematics of the proboscis, we turn to the dynamics of pressure. As a demonstration of pressure increase during eversion, we punctured the body wall using a capillary tube (outer diameter 0.75 mm, inner diameter 0.5 mm). The rise of hemolymph in the tube reflects both capillary pressure, which is a function of the fluid and material properties of the tube, and internal body pressure, which is controlled by the worm’s muscles. According to Jurin’s law [33], the capillary rise *h*_*c*_ is given by *h*_*c*_ ~ 4*σ/ρgd*, where *σ* is the surface tension of haemolymph, *ρ* is the haemolymph density, *g* is gravitational acceleration, and *d* = 0.5 mm is the capillary diameter. When the worm is at rest, the hemolymph rose to 97.5 mm in the capillary, of which 58.8 mm can be attributed to capillary pressure alone using Jurin’s law. The remaining 38.7 mm of head corresponds to the worm having an internal baseline pressure of 0.38 kPa. During eversion, the fluid rose even higher, indicating an increase in internal pressure (Figure 7a).

The pressure probe was inserted approximately 5 cm from the head to avoid puncturing the fang/venom gland complex or buccal tube. We simultaneously recorded internal pressure build-up and filmed the proboscis eversion. Figure 7c shows a representative time course of pressure across 160 seconds. The yellow-highlighted regions indicate periods when the proboscis was everted. In this trial, the peak pressure during eversion reached 2.18 *±* 0.11 kPa, which is comparable to 1.36 *±* 0.56 kPa, the average across 11 eversions. The measured pressure was comparable to the internal pressure in peanut worms (*Sipunculus nudus*, 1.66 kPa,) [34] and the pressure in cactus worms (*Priapulus caudatus*, 2.47 *±* 0.37 kPa) [15]. The average duration of eversion and retraction was 9.3 *±* 5.0 seconds in air. This duration is similar to our previously measured values in worm kinematic experiments, where eversion and retraction in air was 11.9 *±* 5.6 seconds. We notice the eversion pressure and duration have large variability indicating a range in eversion behavior.

The staircase-like pressure pattern (Figure 7c) shows that the internal pressure often builds up close to the eversion threshold without triggering eversion. We surmise that reaching a threshold value of internal pressure is not the sole trigger for eversion. Additional muscular regulation may be involved. For instance, during the first eversion in Figure 7c, pressure first rises to 1 kPa and then, through a series of spikes, rises to the highest value of 2.2 kPa. However, in the second and third eversions, pressure only rises to 0.7 kPa with a single spike in pressure before eversion. We also observed via touch that the worm’s body became noticeably turgid before eversion, indicating a buildup of internal pressure.

Figure 7b shows the time course of internal pressure (blue) and proboscis length (orange) during a single eversion, corresponding to the first eversion in Figure 7c. As pressure builds within the body wall and triggers eversion, fluid is pumped into the proboscis, and we hypothesized a pressure drop after elongation. In the trial shown in Figure 7b, the worm generated its highest internal pressure 1.5 seconds after reaching its maximum eversion length. However, this delay was not statistically significant for the other 14 eversion events, where the time delay was measured to be 0.03 *±* 0.74 seconds.

Figure 7d presents a time sequence of proboscis eversion alongside a schematic illustration. The worm initiates eversion by increasing pressure, extending the proboscis until it reaches its maximum length, indicated by the orange asterisk. In the example shown, the proboscis continues to expand radially, reaching its maximum volume (shown by the blue asterisk). In five other eversion events, peak eversion length and peak proboscis volume typically coincide, with a time delay time difference of 0.16 *±* 0.88 seconds (Figure 7e).

### E Work and factor of safety

Physically, both eversion and retraction are actuated by muscles. But eversion has an intermediary step of using fluid hemolymph to transmit muscular energy to the proboscis. We may idealize the bloodworm as a two-piston apparatus shown in Figure 1b. As the body wall contracts, pressure builds in the fluid. Work by the body wall muscles, *W* = *FdL* equals the integral of the product of muscular force *F* and average displacement of the muscles *L*. Simply applied pressure alone is not enough to do work; there must also be displacement in the form of the eversion of the proboscis. The force on the proboscis is *F* = *PA* where *P* is the internal body pressure and *A* is the cross-sectional area of the proboscis. We may write the work as *W* = *PdV* where *dV* = *AdL*. Since we cannot accurately measure the pressure and volume change simultaneously, we estimate the work done as *W*_*p*_ = *P*_*eversion*_Δ*V*, where Δ*V* = 785*mm*^3^ is the volume of the proboscis. Using the peak pressure *P*_*eversion*_ = 1.36 kPa, the work of eversion in air is 1.1 mJ, which is equivalent to 0.0003 calories. Work can also go towards the energy of bending the tissue and viscous dissipation in the hemolymph. From the math section and measured material properties, we show the work required for bending *W*_*b*_ and viscous dissipation *W*_*v*_ is negligible, with values of *W*_*b*_ = 5.6 *×* 10^−7^*mJ* and *W*_*v*_ = 8.0 *×* 10^−4^*mJ*, respectively. Using calorimetry, Arrazola-Vasquez *et al* showed that earthworms spend 10-30 J per day burrowing [35]. If bloodworms have similar energy budgets, this would allow for approximately 9,000 to 28,000 full eversions per day.

We next calculate the safety factor of eversion, defined as the ratio of the pressure during eversion to the maximum pressure allowed by material properties. Using the fracture stress obtained from material testing, we calculate the fracture pressure *P*_*frac*_ using Equation 2, yielding *P*_*frac*_ = 67 kPa. Given the eversion pressure *P*_*eversion*_ = 1.36 kPa measured from the pressure experiments, this results in a safety factor of approximately SF_*stretch*_= 49. For comparison, the ASME Boiler and Pressure Vessel Code typically requires safety factors in the range of 3.5 to 4 for pressurized systems [36].

We next calculate the safety factor SF_*bend*_ due to bending of the proboscis. The greatest strain of eversion is highly localized: specifically, it occurs at the buccal tube’s inflection points, which we show using a CAD model in Figure 5e. Here, the red surface indicates the external buccal tube lining that becomes internal when everted; similarly the pink internal tube becomes the outside when the tube is everted. As the inflection points move down the buccal tube, more of the tube is inverted in a process similar to rolling up a shirt sleeve. Assuming the worst possible case for the strain, the proboscis skin bends 180 degrees with a strain of *ϵ*_*bend*_ = 1. Given the failure strain of *ϵ*_*max*_ = 3, the safety factor for bending is SF_*bend*_ = 3. The proboscis is clearly adapted for eversion. We note the body wall would not be a suitable material for eversion since it has a failure strain of 0.759.

**Fig. 5:**
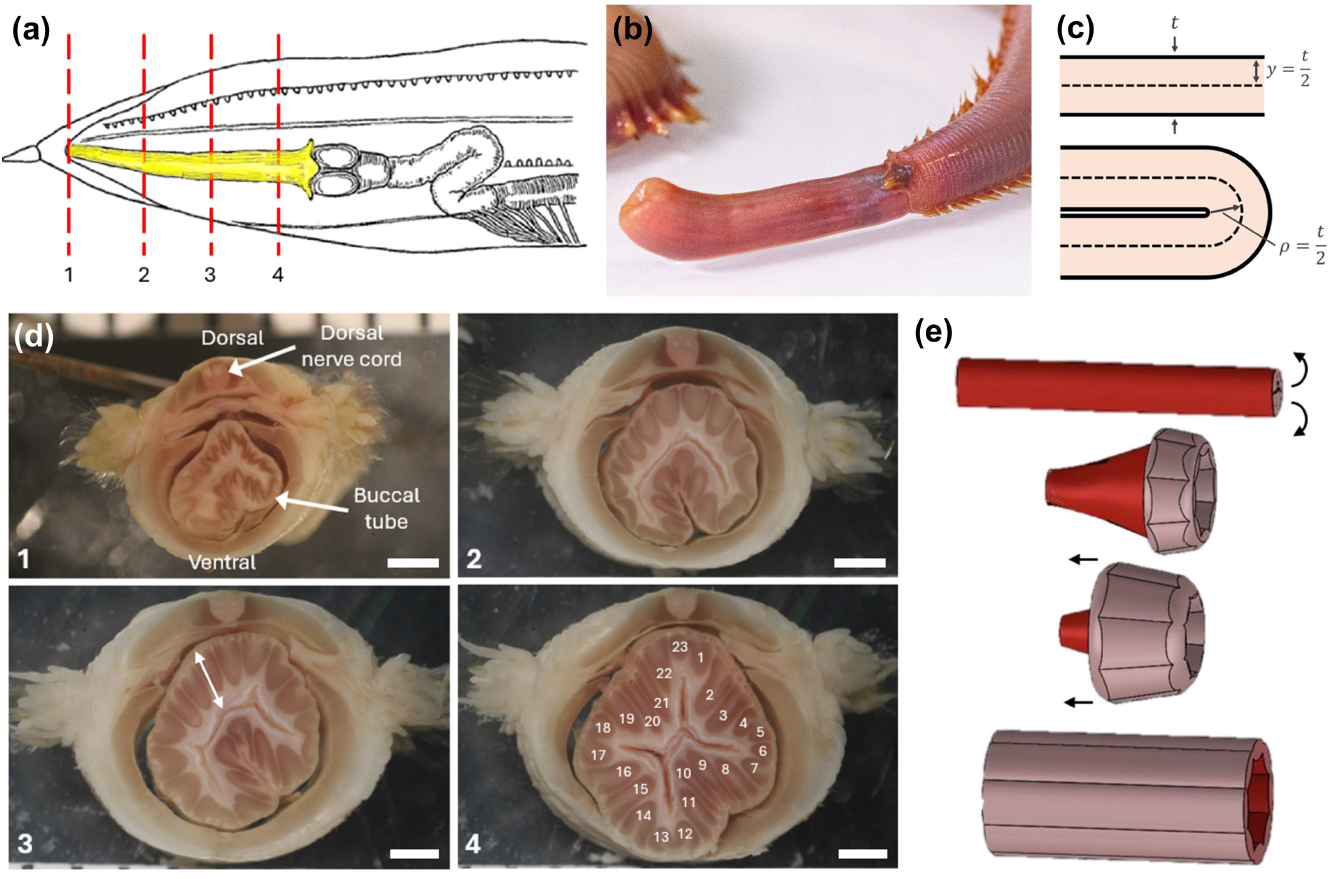
Everting structure of bloodworm buccal tube. (a) Schematic diagram of the bloodworm with highlighted buccal tube (yellow) and marked positions for histological analyses, adapted from Wells (1937) [24]. (b) An everted buccal tube showing longitudinal stripes indicative of longitudinal muscles. (c) Schematic of the sagittal view of bloodworm proboscis in straight and folded state. Transverse histological sections at the longitudinal positions marked in part a. The proboscis thickness is measured using the arrow given in d3. The fourth image shows numbering of 23 individual longitudinal muscles. Scale bars, 1 mm. (e) Schematic of the buccal tube during eversion, where red represents the internal surface in the everted state and pink represents the inner surface that becomes exposed upon eversion.

**Fig. 6:**
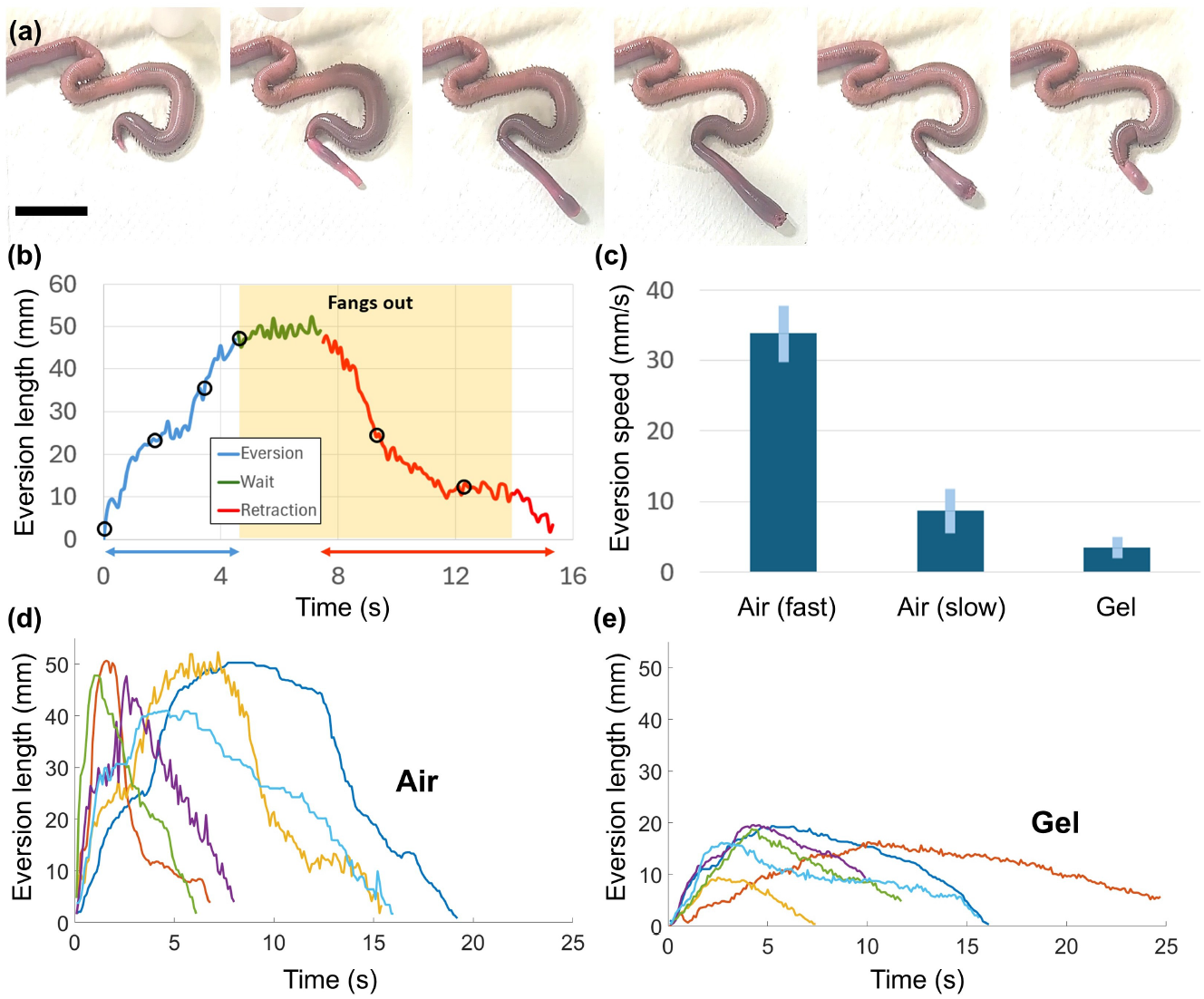
Eversion dynamics of bloodworms in different environments. (a) Sequential images of a bloodworm undergoing proboscis eversion in air. (b) Time course of eversion length, highlighting different phases: eversion (blue), waiting (green), and retraction (red). Hollow circles correspond to the time points of the images in (a). The shaded yellow region marks the phase where the fangs are visible. (c) Comparison of eversion speeds in different environments. Error bars represent standard deviation. (d-e) Eversion length over time in air (d) and gel (e), illustrating faster and more complete eversion in air compared to gel.

**Fig. 7:**
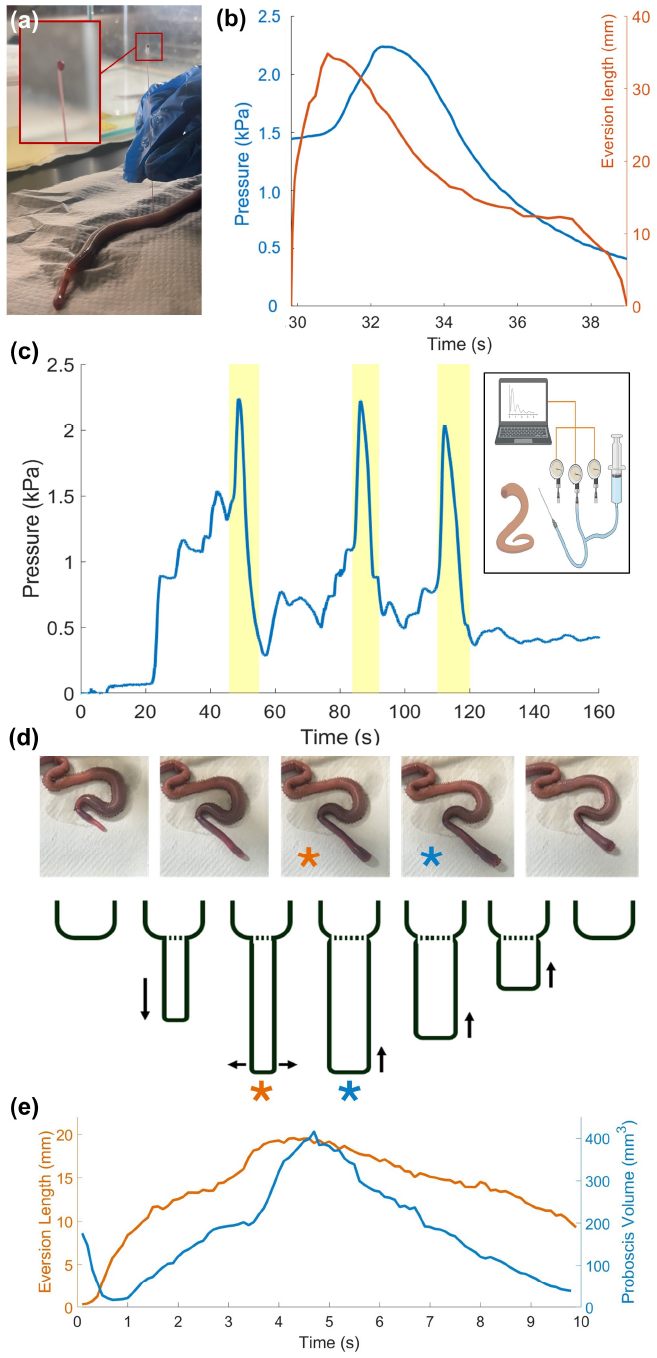
Internal pressure measurement during bloodworm eversion. (a) A capillary tube is inserted into the body of a bloodworm to monitor internal pressure. Upon eversion, blood rises visibly within the tube, with the inset showing overflow at the tube tip. (b) Simultaneous time course of internal pressure (blue) and eversion length (orange) in a live bloodworm. (c) Repeated proboscis eversion and retraction are captured in the pressure graph. Time *t* = 0 denotes needle insertion. Yellow-highlighted regions indicate periods of visible eversion, with the first event corresponding to the data in (b). The inset shows schematic of the experimental pressure measurement setup (created in BioRender). (d) Sequential images of bloodworm along with illustrations of everting proboscis. Time course of eversion length (blue) and eversion volume (red). The blue dashed line and star indicate the peak eversion length, while the red dashed line and star indicate the peak proboscis volume.

In summary, we present two safety factors, one based on stretching and the other based on bending:

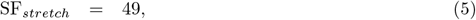

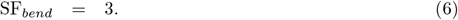

We conclude that the worm is in greater danger of material failure due to bending then stretching.

### F Comparative analysis of eversion

Through literature and online video search, we conducted a comparative analysis of nine species of everting organisms including invertebrates (ribbon worm *Gorgonorhynchus repens*, cactus worm *Priapulus caudatus*, caterpillar *Papilio xuthus*, snail *Cornu aspersum*, sunbeam caterpillar *Curetis acuta*) and vertebrates (frog *Lithobates catesbeianus*, shark *Galeoverdo cuvier*, ray *Tetronarce nobliana*). We measured eversion speeds and the sizes of everting organs from online videos and assumed the animals had adult body masses, whose values we found from previous literature (we provide videos in the open-source repository [37]). For comparison, we also include two soft robots: (1) vine robot for navigation through constrained environments [4] (Figure 8i), and (2) growing sling to support and move individuals [5] (Figure 8j) but we do not include the robots in the trendline calculations. Before we report the regime diagrams, we briefly review the context in which these animals evert.

**Fig. 8:**
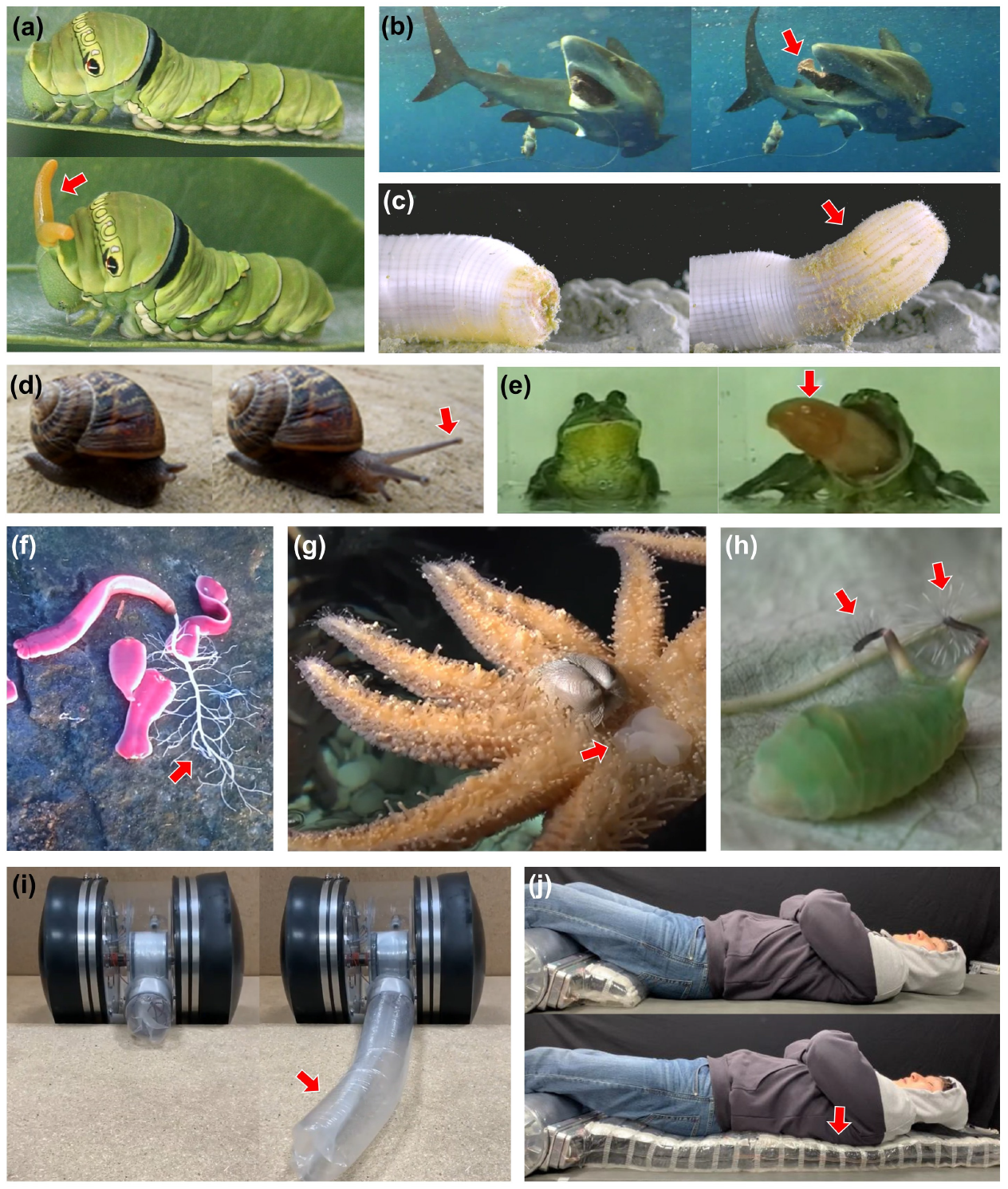
Before and after images of everting animals and robots, with red arrows indicating the everting structures. (a) The caterpillar, *Papilio xuthus*, which uses its tentacle as a defensive eversible organ (image source: Wikipedia, 2011). (b) A shark displaying stomach eversion, a behavior used to expel indigestible materials (image source: YouTube, Jeff Stein). (c) A cactus worm everting a structure called the praesoma as part of their feeding (image source: YouTube, The Naturalist Project). (d) A snail showing its tentacle extension (image source: YouTube, Musselshell). (e) A frog utilizing its eversion mechanism to catch prey with its tongue (image source: Fuji TV, “Spring of Trivia”, 2003). (f) A ribbon worm, *Gorgonorhynchus repens* (probable) exhibits densely branching structure of proboscis (image source: YouTube, Katsukaida Iori). (g) Sunflower sea star, *Pycnopodia helianthoides* possesses stomach that can extend outside the mouth to digest prey (image source: Vancouver Aquarium). (h) Angled sunbeam caterpillar, *Curetis acuta*, everts and oscillates tentacle organs located in the two periscopes (image source: Youtube, rarasukekun) (i) Vine robots demonstrating robotic eversion [78]. (j) A growing sling designed for patient transfer [5].

For slow-moving marine worms and caterpillars, eversion provides speed for attack or defense [24, 25, 38–43]. Ribbon worms *Gorgonorhynchus repens* evert a highly branched proboscis, likely to increase the contact area with which to secrete toxins to their prey (Figure 8f) [44]. Phylum Priapulida evert a structure called the praesoma as part of their feeding (Figure 8c). Hunter and Elder (1989) investigated praesoma eversion in the cactus worm *Priapulus caudatus* [15]. Caterpillars of swallowtail butterflies, family Papilionidae, evert a pair of defensive appendages called osmeterium when threatened [40]. Osmoterium can release toxins to deter attackers or honeydew-like secretions that facilitate their mutualistic relationship with ants [45]. Angled sunbeam caterpillar *Curetis acuta* larvae have “periscopes” on their rear end that evert toxin-tinged dandelion-like appendages that quickly rotate to ward off potential threats (Figure 8h)

Snails have two sets of eversible head tentacles tipped with eyestalks or olfactory organs (Figure 8d) [46]. These are called tentacles rather than limbs because they are long and flexible and have an ability to retract. A tentacle contains three string muscles and a main tentacle retractor muscle. These muscles control retraction and directional control of the tentacle [47]. The anterior tentacles are dense in chemoreceptors, which makes its olfaction comparable to that of vertebrates [48].

Organisms may evert their digestive systems for a variety of reasons. When consuming molluscs, the starfish *Pycnopodia helianthoides* pries the mollusk partially open and then everts its stomachs between the shells (Figure 8g). Animals that cannot vomit use eversion to remove unwanted contaminants or debris. For example, sharks and frogs evert their stomachs through their mouth [49, 50] (Figure 8b). This is medically termed gastric prolapse, and it is analogous to vomiting in mammals [51]. Stomach eversion in sharks is observed during fishing due to irritation of the shark’s digestive system from an ingested fish hook [49]. This observation dates back to at least 300 years ago when fishermen caught sharks with their stomachs protruding out of their mouths [51]. Stingrays evert their large intestine through their cloaca in response to stress or handling [52]. Sea cucumbers *Bohadschia argus* also exhibit a dramatic defensive behavior, known as evisceration, where internal organs such as the gut, respiratory tree, and Cuvierian tubules are expelled through the cloaca. However, this involves rupture and expulsion without the structure turning inside out, so they were excluded from our scaling analysis [53].

In frogs (Figure 8e), eversion can occur if the frog suffers from a terminal disease or gastric prolapse [54]. Frogs have a mucus layer in their stomachs that is thinner than those of mammals which may make their stomachs easily irritated. Frog stomachs are regulated by endocrine cells. When the endocrine balance is disrupted, the frog’s stomach contracts and everts out of its mouth [55].

We next turn to the scaling of eversion, beginning with pressure generated. Figure 9a shows a regime diagram comparing the magnitude of pressures across animals. Internal pressure is indicated by the black markers, suction pressure is indicated by blue markers, and eversion pressure is indicated by red markers. Body masses vary by ten orders of magnitude while pressures only vary across 2-3 orders of magnitude. This near constancy of pressure is due to the fact that pressure is generated by muscular stress which is physiologically limited by the strength of muscle tissue [56]. Internal pressures are generally positive either because the heart is pumping blood or body wall muscles are contracting to circulate hemolymph to deliver oxygen. The internal pressure ranges from 0.5 to 12 kPa. Invertebrates such as earthworms, snails, and crabs have low internal blood pressures of 1–2 kPa [57–59], which are similar to the eversion pressure of bloodworms, cactus worms, and peanut worms [15, 34]. Squid are quite athletic and they have high internal pressure of 5.5 kPa compared to other soft-bodied invertebrates [60], while humans maintain an even higher 12 kPa in their circulatory system. Suction pressures are remarkably constant across scale, remaining near 10 kPa for mosquitoes, humans, and elephants [61–63], despite spanning nine orders of magnitude in mass. Eversion pressures are within the same range as these other pressures, suggesting that everting animals satisfy the same biological limits for pressure as other animals.

**Fig. 9:**
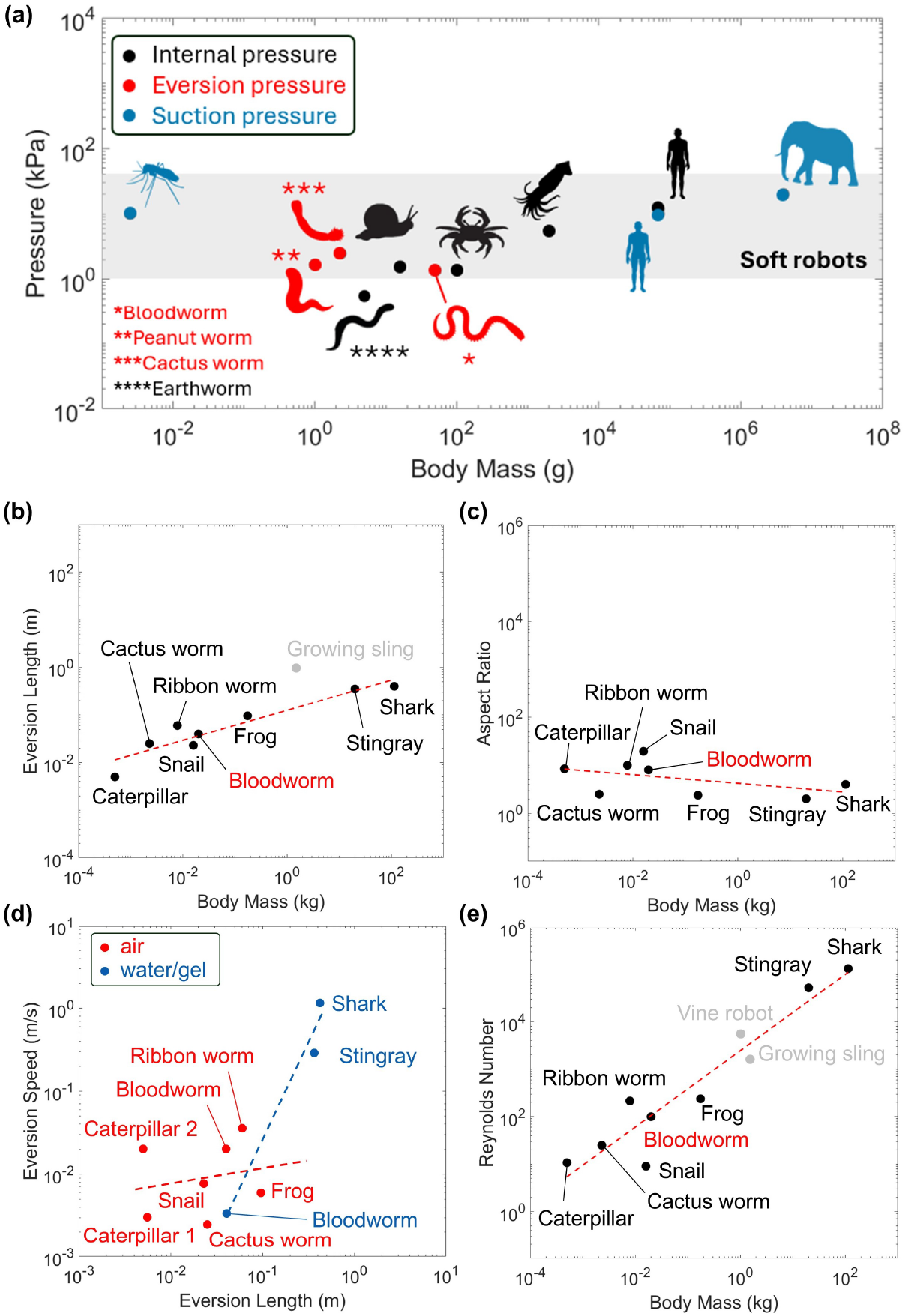
Regime diagrams and scaling across various animals. (a) Regime diagram comparing internal, eversion, and suction pressure of soft bodied animals, mosquitoes, humans, and elephants (silhouette images from Shutterstock, Istockphoto, Onlygfx, Pixabay, and Vexels). (b) Scaling relationship between eversion length and body mass for everting animals and robots. The linear fitting is given as a red dashed line. (c) Scaling relationship between Reynolds number and body mass. (d) Scaling relationship between aspect ratio and body mass. (e) Scaling relationship between eversion speed and eversion length with best-fit lines. Red markers indicate animals everting in air (slope = 0.18), while blue markers represent animals everting in water or gel (slope = 2.34).

Figure 9b shows the relationship between everting organ length *L* and body mass *M*. The best fit line may be written:

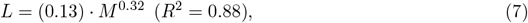

where *M* is in kg and *L* is in m. The length *L* has a power exponent of 0.32, or near isometry (*M* ^0.33^), meaning that the proportions of the animal are constant.

Many everting organs can be modeled as cylinders with diameter *D*. For the rectangular cross-section of the growing sling robot [5], we calculated a hydraulic diameter based on its area and perimeter. Figure 9c shows the everting organ aspect ratio, We defined an aspect ratio, or length/diameter,

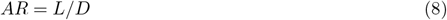

where the best fit

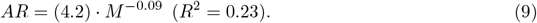

indicates aspect ratio is ranges between 2 and 20. Note that the scaling for aspect ratio shows a low *R*^2^ value and the near-zero slope, suggesting no clear dependence of aspect ratio on body mass. We thus see that everting organs satisfy a number of purposes and they can have a broad range of proportions. Snails, which evert their long tentacles, have the longest aspect ratio, while the sharks, stingrays, and frogs also evert their intestines and stomachs, which are squat-shaped in comparison.

Figure 9d shows the relationship between eversion speed *U* and everting organ length *L*. Since speed depends on the media, the animals were grouped into two categories, with those that evert in air in red and the others that evert in water or gelatin in blue, with best fits given by

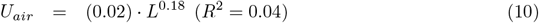

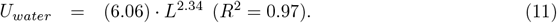

where *L* is in m and *U* is in m/s.

Darcy Thompson reported a “Froude’s Law of Equivalence of Velocities” which relates the speed and size of an organism: *U* ~ *L*^0.5^ [64]. It was inspired by the motion of boats on water, in which the wave speed of surface gravitational waves is 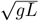 where *L* is the wavelength. Boats that traveled at the same Froude number, Fr= *U* ^2^*/*(*gL*) would be dynamically similar, with boats traveling at high Fr rocking up and down because they are climbing past their own bow wave. Other workers found that Froude’s Law was descriptive of bird flight [65–67], fish swimming [68], and even water-walking insects [69]. For birds, Froude’s law holds because the bird must support its body weight *Mg* with the drag force on its wings which scales as *ρU* ^2^*A*^2^ where *U* is its wing velocity, *A* is the wing area, and *ρ* is the density of the air. Thus for weight support, *Mg/ρU* ^2^*A*^2^ ~ *L*^3^*/*(*U* ^2^*L*^2^), and thus *U* ~ *L*^0.5^.

Everting speed in air showed a weak dependency on length with the power exponent of 0.18 which is lower than Froude’s law (0.5), and the fit had a relatively low *R*^2^ value. In contrast, everting animals in water followed a markedly different trend, with *U* ~ *L*^2.34^. As expected, bloodworm proboscis eversion is faster in air than in water. But surprisingly, sharks and rays everting in water had the highest speeds of all the animals studied. For comparison, swimming speeds scale as *U* ~ *L*^0.43^ in aquatic vertebrates including fishes, mammals, and penguins [70], and as *U* ~ *L*^0.3^ for squid [71], which use contraction of muscles to jet water similar to the use of muscles to squeeze fluid into an everting organ.

What causes the speed-scaling for everting animals? The driving force of eversion is the contraction of circumferential muscles. The muscle force scales as the cross-sectional area of the body as *F* ~ *σL*^2^ where *L* is the length scale and *σ* is the muscle stress. Assuming the same type and density of muscle fibers, their stress output is nearly constant across species [65]. Thus we have

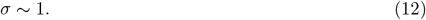

When the muscle contracts and decreases the radius of the cylindrical body, we consider the hoop stress from Equation 3, *σh* ~ *RP*. The *h* and *R* cancel as they can be scaled as *L*. Thus we have

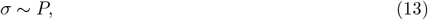

where *P* is the required pressure for eversion, and can be scaled as dynamic pressure of moving fluid as *P* ~ 1*/*2*ρU* ^2^ at an eversion speed *U*, assuming that the eversion speed is comparable to the fluid velocity within the body. Combining with Equations 12 and 13, we have the eversion speed is constant: *U* ~ *M* ^0^ ~ *L*^0^, which compares poorly with both eversion in air *U* ~ *L*^0.18^ and eversion in water *U* ~ *L*^2.34^. Moreover, our model does not explain the surprisingly higher speeds for aquatic eversion. It is possible that aquatic animals may use hydrodynamic forces to increase their eversion speed. For example, sharks may swim forward while everting, using the drag force on their stomach to pull it out.

Lastly, Figure 9e shows the Reynolds number

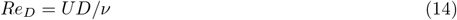

where *U* is the eversion speed, *D* is the diameter of the everting body and *ν* is the kinematic viscosity of the internal fluid of the everting body, where we assume the viscosity of water for animals and the viscosity of air for robots. The best fit is

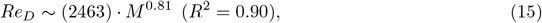

where robots are not included in the scaling. Notably, several animals such as rays and sharks exhibit eversion Reynolds numbers exceeding 1000, approaching the transitional regime to turbulence. This stands in contrast to most biological systems, where internal flow, such as in cardiovascular and olfactory pathways, typically operates at low Reynolds and Womersley numbers (the ratio of inertia to viscosity for oscillating flows). For instance, blood flow in humans is characterized by a Reynolds number of 4,000 [72], and Womersley numbers across species, from pygmy marmosets to deer spanning nearly three orders of magnitude in mass, remain within a narrow range of 0.2 to 2.5 [73]. This feat is accomplished by the use of branching pipes which help prevent the flow of blood or air from transitioning to turbulence.

## V Discussion

The bloodworm offers a biological analog that achieves robust eversion and coordinated retraction through the interplay of fluid pressure and retractor muscles. Pressure measurements suggest that internal pressure alone does not trigger eversion; bloodworms are capable of resisting eversion even under elevated pressure. This behavior resembles latch mechanisms observed in many biological systems [74]. Eversible structures paired with retractor muscles are found across diverse species, including bloodworms and cactus worms. Although these species belong to different phyla (Annelida and Priapulida), they have independently evolved similar strategies for feeding and locomotion via eversion. Interestingly, the alignment of retractor muscles differs: bloodworms exhibit asymmetric arrangements, whereas cactus worms display symmetric organization [75]. This functional decoupling between internal fluid pressure and retractor muscle contrasts with most soft robotic systems, where eversion and pressure are simultaneously modulated. Independent control over these two actuation modes in a biological system offers a model for actuator design in soft robotics.

The bloodworm compactly stores its everting organ when not in use. In the resting state, the deflated buccal tube and intestine are buckled to reduce their length, and the retractor muscle displays pronounced wrinkles and folds, potentially functioning as a form of self-organized origami [28]. These natural strategies may provide insights into repeatable retraction. Recent work has explored material scrunching mechanisms to improve retractability in soft-growing robots [76], highlighting the relevance of biologically inspired folding and compaction strategies.

While this study focused on linear eversion, we also observed bloodworms directing their proboscis along curved paths. Such steering likely involves localized muscle activation, and investigating these dynamics may uncover more complex control strategies. We only observed two speeds of eversion, a fast and a slow. Further experiments involving variations in substrate stiffness or surrounding fluid resistance could help elucidate whether bloodworms evert using feedback from the environment.

## VI Conclusion

Bloodworms achieve rapid eversion by increasing internal fluid pressure which extends and expands the proboscis. The wrinkled retractor muscles and buckled configuration of internal organs show the worm’s compact storage and deployable design. Material testing also revealed a safety factor of 49 against linear stretch and a safety factor of 3 with respect to bending failure. The bloodworm proboscis presents an example of a natural hydraulic actuator with implications for miniaturized, extendable, and reversible robotic systems.

## VII Acknowledgements

Funding is provided by NSF grant 2344314 and 1755314 (JTT). We thank Stanislav Y. Emelianov for hosting ultrasound experiments, Brandon Dixon for hosting pressure experiments, and Lakshminarayanan Mahadevan for suggesting the regime diagram. We are also grateful to Anna Alvazez and Elliot Hawkes for insightful discussions on vine robot applications.

## Suppementary material captions

- Download link: [Google Drive](https://drive.google.com/drive/folders/1SNlIBaq3-v28XySznLGYj43yh3wXS6Eq?usp=sharing)
- Supplementary video 1. Transverse ultrasound scan of a bloodworm with the proboscis in its retracted state.
- Supplementary video 2. Sagittal ultrasound scan of a bloodworm with the everted proboscis.
- Supplementary video 3. Example video of bloodworm eversion in air. The eversion speed is approximately 20 mm/s.
- Supplementary video 4. Example video of bloodworm eversion in gelatin. The eversion speed is approximately 3 mm/s.
- Supplementary video 5. Animal eversion clips corresponding to the regime map.

